# fullsibQTL: an R package for QTL mapping in biparental populations of outcrossing species

**DOI:** 10.1101/2020.12.04.412262

**Authors:** Rodrigo Gazaffi, Rodrigo R. Amadeu, Marcelo Mollinari, João R. B. F. Rosa, Cristiane H. Taniguti, Gabriel R. A. Margarido, Antonio A. F. Garcia

## Abstract

Accurate QTL mapping in outcrossing species requires software programs which consider genetic features of these populations, such as markers with different segregation patterns and different level of information. Although the available mapping procedures to date allow inferring QTL position and effects, they are mostly not based on multilocus genetic maps. Having a QTL analysis based in such maps is crucial since they allow informative markers to propagate their information to less informative intervals of the map. We developed `fullsibQTL`, a novel and freely available `R` package to perform composite interval QTL mapping considering outcrossing populations and markers with different segregation patterns. It allows to estimate QTL position, effects, segregation patterns, and linkage phase with flanking markers. Additionally, several statistical and graphical tools are implemented, for straightforward analysis and interpretations. `fullsibQTL` is an `R` open source package with `C` and `R` source code (GPLv3). It is multiplatform and can be installed from https://github.com/augusto-garcia/fullsibQTL.

## 1 INTRODUCTION

Mapping quantitative trait loci (QTL) unveils the genetic architecture of quantitative traits (*e.g.* yield, fitness) and has several applications, such as breeding and evolutionary studies. In plant breeding, this information can be integrated into breeding programs with marker-assisted selection tools aiming to understand and boost such traits. Interval mapping (Lander and Botstein, 1989), composite interval mapping (Zeng, 1993, 1994), and multiple interval mapping (Kao et al., 1999) are some of the available approaches to perform QTL mapping. In populations developed from inbred (homozygous) lines, such as F2s, backcrosses and recombinant inbred lines, these mapping methods are well established and implemented in a plethora of good software, for example: Broman et al. (2003); Van Ooijen (2004); Wang et al. (2012); Meng et al. (2015). However, for outcrossing species without inbred lines (*e.g.* citrus, eucalyptus, sugarcane, loblolly pine, and rubber tree), there is a lack of freely available software programs for QTL analysis. For biparental populations derived from the cross between non-homozygous parents, it is necessary to estimate QTL number, effects and position, as well their linkage phase with markers, and segregation patterns. This effort relies on a linkage map that provides the phase of markers in both parents and their complete transmission pattern to the offspring, which is usually thoroughly achieved by using multipoint approaches.

`fullsibQTL` is a free R package that performs composite interval mapping in full-sib progeny (F1 population) derived from a bi-parental cross between two non-homozygous parents. It requires a previously estimated genetic map, considering markers with different segregation patterns, as described by (Wu et al., 2002b). It implements the model proposed by Gazaffi et al. (2014), allowing to estimate the number of QTLs, their position, effects, segregation patterns, and linkage phases with flanking markers. This methodology has been successfully applied in different studies (Souza et al., 2013; Margarido et al., 2015; Costa et al., 2016; Balsalobre et al., 2017; da Silva Pereira et al., 2017; Barreto et al., 2018; Conson et al., 2018; Curtolo et al., 2020; Soratto et al., 2020), but currently there is no competing software with such features.

## 2 FEATURES

`fullsibQTL` is a toolbox to perform QTL mapping based on composite interval mapping using the model proposed by Gazaffi et al. (2014) for outcrossing species. It considers that a multipoint genetic map was previously estimated with package `OneMap`, also freely available and widely used (Margarido et al., 2007, 2020). Briefly, the model has three genetics effects, one additive effect for each parent and one for the interaction of these effects (dominance). These parameters are estimated based on the maximum likelihood for mixture models, with the expectation maximization algorithm. QTL genotypes probabilities are calculated using a multipoint approach based on hidden Markov models (Wu et al., 2002a; Mollinari et al., 2009). This allows the estimation using the information of all the markers and individuals, even if there is missing data (Wu et al., 2002a).

The package also has a function to select cofactors using multiple linear regression through stepwise and information criteria (R Core Team, 2017). Permutation tests (Churchill and Doerge, 1994) for threshold determination can be implemented, combined with modifications suggested by Chen and Storey (2006) for a more relaxed threshold. The model also can have other covariates. Moreover, several functions were developed to provide graphical (Fig. 1) and text output. Additionally, interval mapping and inclusive composite interval mapping options (Li et al., 2006) are also included. To illustrate the usage of the package, a simulated dataset from Gazaffi et al. (2014) and its analysis are included in the package, with a full tutorial and vignettes, explaining how to deal with partially informative markers.

**Figure 1.**
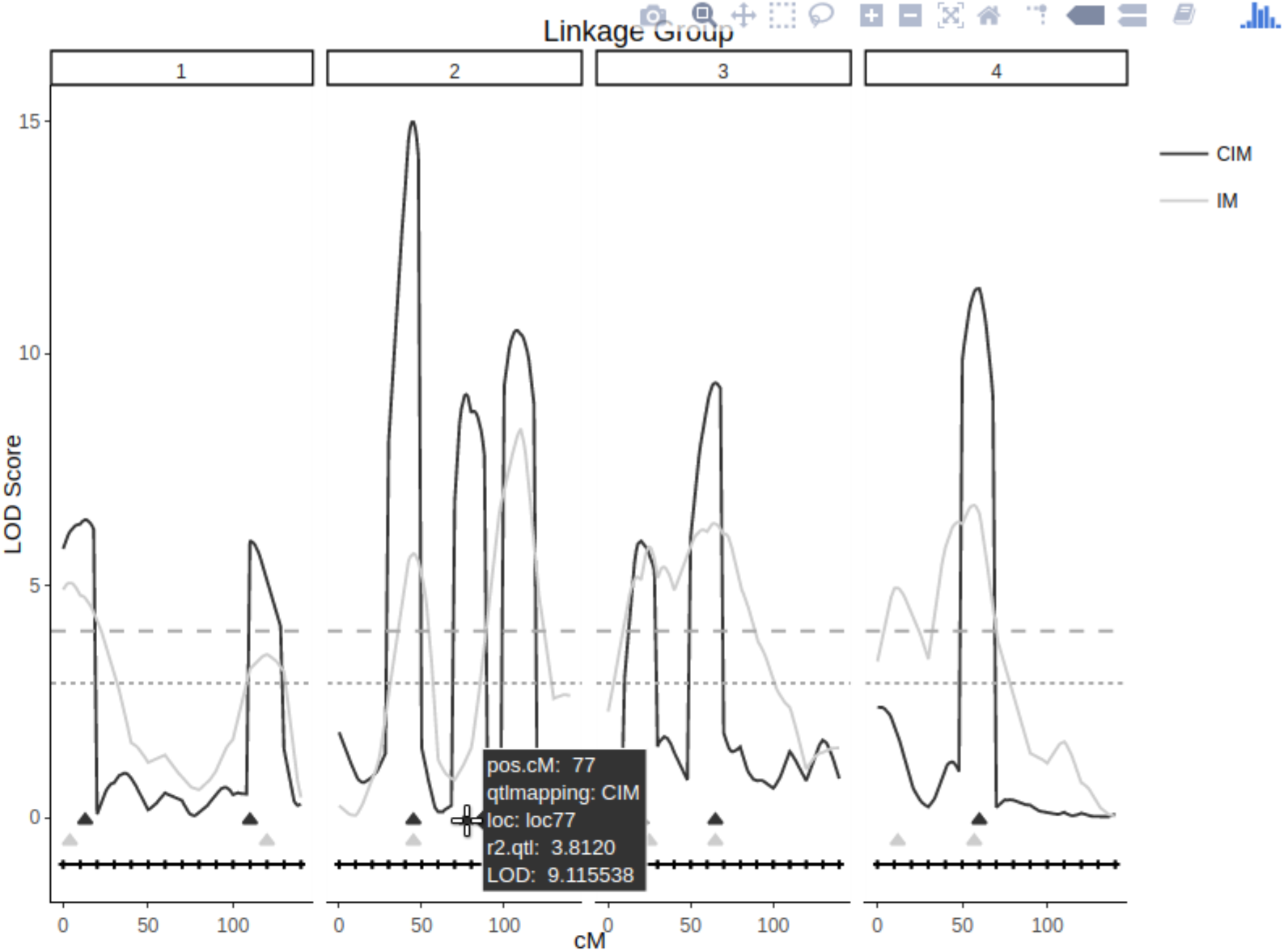
Screenshot of a `fullsibQTL` interactive `HTML+plotly` graphic. It provides a comparison of composite interval mapping and interval mapping approaches for a simulated population. The x-axis shows genetic map for each linkage group, and the y-axis the corresponding logarithm of the odds score for testing the presence of putative QTL. At the top right, there are buttons to zoom, box selection, etc. The mouse hover shows *R*^2^, position, and logarithm of the odds scores.

With the results of QTL analysis, based on logarithm of the odds scores of hypothesis tests (Gazaffi et al., 2014), users can visualize their genetic effects, statistical significance, segregation patterns and linkage phase with flanking markers. Based on a least square approximation, `fullsibQTL` can also compute the amount of phenotypic variation explained by each QTL in the model (the coefficient of determination, *R*^2^).

## 3 CONCLUSION

Based on a sound QTL mapping methodology for outcrossing species, and on a reliable multipoint genetic map, provides a QTL mapping toolbox to study the genetic architecture of quantitative traits in outcrossing species. It allows access to relevant information, and includes several graphical and analytical features, providing an ideal environment for interaction with users. Its initial versions were extensively tested and successfully applied several real world problems.

## ACKNOWLEDGEMENTS

Authors thank André Conson and Marianella Quezada for their helpfull feedbacks as beta testers, and Karl Broman for sharing code.

